# Intracellular binding pocket revealed in the human bitter taste receptor TAS2R14

**DOI:** 10.1101/2024.04.10.588278

**Authors:** Lior Peri, Donna Matzov, Dominic R. Huxley, Alon Rainish, Fabrizio Fierro, Liel Sapir, Tara Pfeiffer, Lukas Waterloo, Harald Hübner, Dorothee Weikert, Yoav Peleg, Peter Gmeiner, Peter J. McCormick, Masha Y. Niv, Moran Shalev-Benami

## Abstract

Bitter taste receptors (TAS2Rs), a subfamily of G-protein coupled receptors (GPCRs) expressed orally and extraorally, elicit signaling in response to a large set of ligands. Among the 25 functional TAS2Rs encoded in the human genome, TAS2R14 is the most promiscuous, and responds to hundreds of chemically diverse agonists. Here, we present the cryo–electron microscopy (cryo-EM) structure of the human TAS2R14 (hTAS2R14) in complex with its cognate signaling partner gustducin, and bound to flufenamic acid (FFA), a clinically approved nonsteroidal anti-inflammatory drug. The structure reveals an unusual binding mode for FFA, where two copies are bound at distinct binding pockets: one at the canonical GPCR site within the trans-membrane bundle, and the other in the intracellular facet, bridging the receptor with gustducin. Combined with site-directed mutagenesis and the design of a fluorescent FFA derivative for pocket-specific ligand binding BRET assays, our studies support a dual binding mode for FFA in TAS2R14. These results fill a gap in the understanding of bitter taste signaling and provide tools for guided design of TAS2R-targeted compounds.

## Introduction

Bitterness is elicited by the taste 2 receptor (TAS2R, class T) family. In humans, 25 functional TAS2Rs are encoded in the genome (*1*), that are typically classified as class A like GPCRs. However, TAS2Rs lack most Family A hallmark motifs, and share low sequence similarity both within the subfamily members and with other GPCRs (*2*). TAS2Rs respond to a diverse array of tastants, and while some family members are narrowly tuned for specific ligands, others respond to agonists with remarkable chemical diversity (*1, 3, 4*). TAS2R14 is a notable taste receptor that responds to the largest number of agonists, including vitamins (*5*), flavonoids (*6*), pharmaceuticals (*7, 8*) and odorants (*9*). TAS2R14 is abundant in the taste buds but is also expressed in extraoral tissues such as the respiratory (*10, 11*), cardiovascular (*12, 13*) and digestive (*14*) systems. Polymorphisms in the TAS2R14 gene were associated with cardiac physiology (*13*) and male infertility (*15*). TAS2R14 overexpression was also documented in several types of cancer (*12, 16*), and its stimulation was shown to elicit anti-proliferative and pro-apoptotic effects in multiple cancer cell lines (*17–19*). In addition, individuals diagnosed with pancreatic adenocarcinoma that have elevated TAS2R14 expression levels show an extended overall survival compared to those with lower expression of the receptor (*19*). These findings highlight TAS2R14 as a broadly-tuned receptor that responds to a large variety of ligands, and underscore potential therapeutic applications that go beyond bitter sensing.

Upon activation of TAS2Rs in taste cells, intracellular signaling is mediated through coupling to a specialized Gα protein called gustducin, that is primarily expressed in mammalian oral and intestinal tissues to mediate bitter, sweet and umami taste transduction (*20*).

A recent cryo-EM structure of TAS2R46 bound to an agonist and a miniGα gustducin mimetic provided a snapshot of an active bitter taste receptor (*21*). However, due to the large sequence variability of TAS2Rs and their highly varying ligand binding capacity, aspects regarding the mechanism of activation and ligand selectivity in this receptor family remain opaque.

Here we determined the structure of the active human TAS2R14 (hTAS2R14) in complex with miniG-gustducin and bound to flufenamic acid (FFA). FFA is a prescription nonsteroidal anti-inflammatory drug that acts via inhibition of cyclooxygenase-2, and is also one of the most potent TAS2R14 agonists (*8, 22*). The structure reveals a dual binding mode of FFA to the receptor, where two copies of the agonist are bound at distinct receptor loci: one corresponds to the canonical binding site, located at the extracellular part of the receptor, and another is bound at the intracellular side, where it maintains interactions with TMs 3,5 and 7 and is also in direct contact with the alpha-5 helix of G-alpha gustducin. Through mutagenesis studies and the design of a fluorescent FFA conjugate that, along with receptor labeling at either the extra- or intra-cellular sites allowed for the direct detection of the ligand’s binding through Bioluminescence Resonance Energy Transfer (BRET), we further demonstrate the unusual dual binding modes of the ligand and highlight the intracellular pocket as a potential target for drug design.

## Results and discussion

### Overall description of the TAS2R14 constructs used for the structural studies

Full length hTAS2R14 was cloned into a baculovirus vector containing an N-terminal haemagglutinin (HA) signal peptide followed by a cleavable FLAG tag and the first 45 amino acids of rat somatostatin receptor 3 (SSTR3), that were previously shown to facilitate surface expression of TAS2Rs in mammalian cells (*7*). For complex formation, we initially used a full-length human Gα-gustducin (Gα_gus_), of which an ScFV16 epitope was introduced to the N-terminus by replacing the first 15 residues of wild-type gustducin with the corresponding amino-acids of Gαi_1_ (iN15). ScFV16 was shown to stabilize G protein heterotrimer formation in multiple complexes used for structural studies of GPCRs and is thus considered as a universal modulator for the heterotrimeric complex formation (*23–26*). The receptor, Gα_gus_-iN15 and Gβγ were co-expressed in insect cells and purified through tandem affinity steps in the presence of the TAS2R14 agonist FFA and ScFV16. However, despite the formation of a complex, the resulting assembly was highly unstable, and unsuitable for the cryo-EM studies.

We thus further created a miniGα-gustducin (miniG_gus_) construct. The design was inspired by a previously described mini-G_i_ (*27*), where the alpha-helical domain was replaced by an eight residue linker, and seven mutations were added to confer stable protein expression. To test construct activity, we used a split nanoluciferase assay, where the intracellular part of the receptor was fused to a large-BiT (LgBiT) segment, and the C-terminus of a soluble miniG_gus_ was coupled to a small-BiT (SmBiT) fragment (*24, 28, 29*). The proteins were overexpressed in mammalian cell lines, and signaling assays were performed in the presence of FFA at serial concentrations (**Fig. 1G**). The assay indicated a dose dependent response to FFA (EC_50_=46.26 ± 10μM). To confirm the functionality of our miniGα_gus_ we also compared G-protein activity using TRUPATH (**Fig. 1H**), an optimized open-source bioluminescence resonance energy transfer (BRET) based system (*30*). In brief, we used a Gα gustducin with a Renilla luciferase variant (RLuc8) inserted within the alpha-helical region along with an unlabelled beta subunit and a gamma subunit with GFP2 inserted at the N-terminus. This BRET-based system has been calibrated for optimal detection of agonist-induced heterotrimer dissociation (*30, 31*). Despite the difference in design of the complementation and the BRET dissociation assays, we obtained similar concentration response curves (**Fig. 1G** and **Fig. 1H**).

**Fig. 1.**
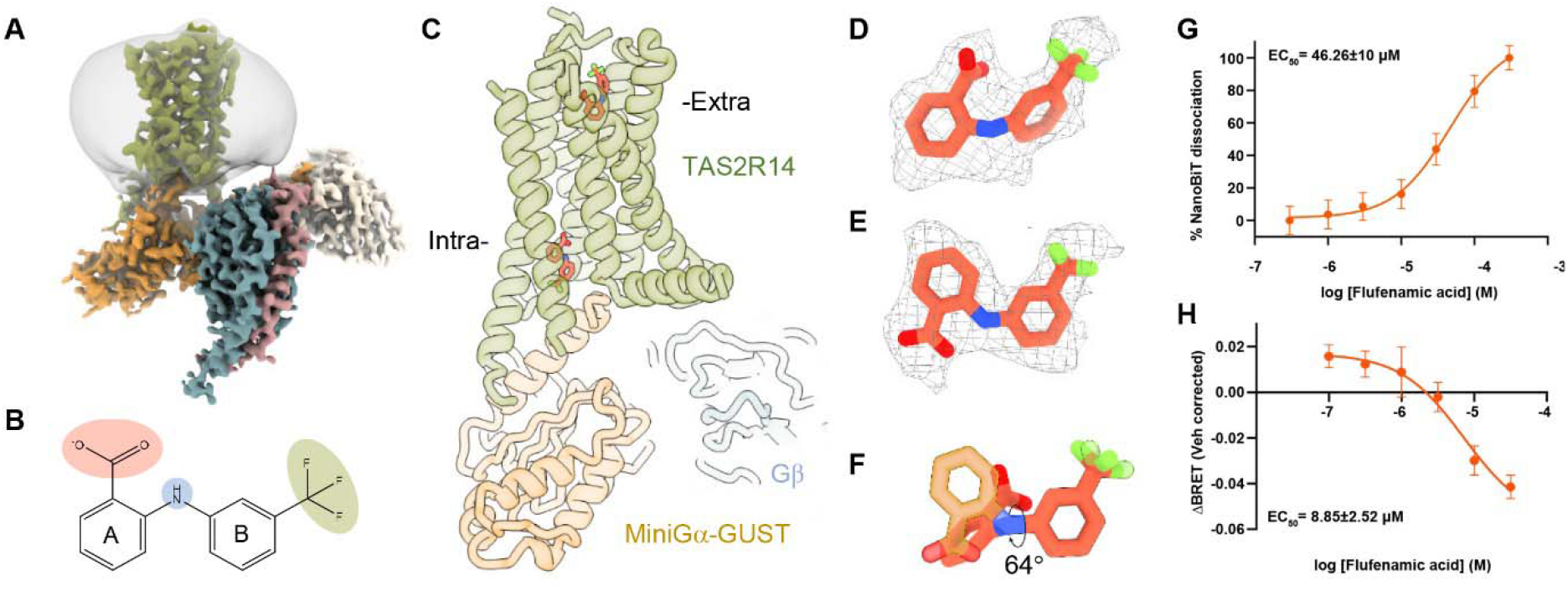
An overall description of the TAS2R14 complex. (**A**) Cryo-EM map of hTAS2R14 bound to the gustducin heterotrimer. TAS2R14 is colored green, gustducin in gold, the beta-gamma subunits are in teal and pink, respectively, and ScFV16 in white. (**B**) The chemical structure of FFA. The ring withr the carboxylic moiety (red) is denoted A, and the ring with the trifluoromethyl moiety (green) is denoted B. The amino linker is highlighted in blue. (**C**) Cartoon representation of the TAS2R14 complex with FFA bound at the canonical (extracellular, extra-) and intracellular (intra-) sites. Colors are as in **A**. (**D**) FFA docked in density for the canonical site. (**E**) FFA in density for the intracellular site. (**F**) Superposition of the canonical (orange) *vs*. the intracellular (gold) conformations of FFA indicating a 64° rotation of ring A relative to ring B. (**G**) Dose response curve for FFA in the mini-gustducin complementation assay designed for this study. Response is measured as nanoBiT dissociation normalized by basal activity of the wild-type receptor to FFA. Data are represented as mean ± SEM from three independent experiments performed in triplicate, indicating EC_50_ value of 46.26 ± 10 μM. (**H**) G-protein dissociation of TAS2R14 in response to FFA in HEK293 cells transiently co-expressing the wild-type receptor and TRUPATH biosensors for the full-length gustducin G protein. Data are represented as mean ± SEM from seven independent experiments performed in triplicate, indicating EC_50_ value of 8.85μM ± 2.52 μM.

For the cryo-EM studies, we used a modified version of the mammalian constructs that also included the beta-gamma subunits. Here, an enhanced affinity SmBiT (HiBiT) (*32*) was fused to the N-terminus of the beta subunit and the gamma subunit was fused to the N-terminus of miniGα_gus_-iN15, as previously described for a G_q_ complex (*25*). Complex formation was induced by supplementing micromolar concentrations of FFA to insect cells co-expressing the plasmids. We then used single-particle cryo-EM to obtain a three-dimensional reconstruction of the hTAS2R14-miniG_gus_βγ complex at a nominal resolution of 2.7 Å. The map revealed the architecture of all complex components, as well as the mode of receptor association with gustducin and the agonist binding pose at the canonical binding site (**Fig. 1D**). Unexpectedly, we found an additional copy of the agonist bound at a distinct pocket facing the intracellular receptor interface (**Fig. 1C,E**). The ligand directly interacts with both the αN5 of gustducin and helices 3, 5 and 7 of the receptor.

### Description of FFA in the canonical binding pocket

TAS2R14 is activated by a large array of chemically diverse ligands (*4, 22, 33*). FFA is among the most potent reported TAS2R14 agonists, and activates the receptor at micromolar concentrations (**Fig. 1G-H**) (*22*). Chemically, FFA consists of two aromatic rings, often referred to as ring A and ring B, that are modified with carboxylic acid and trifluoromethyl moieties, respectively (**Fig. 1B**).

In the cryo-EM structure, FFA engages the canonical binding pocket with the trifluoromethyl moiety pointing towards the extracellular milieu, where it faces a hydrophobic cavity composed of F76^ECL1^, F82^3.25^, F172^ECL2^ and I262^7.34^ (numbers in subscript correspond to Ballesteros–Weinstein numbering (*34*); **Fig. 2A-B**). To test the contribution of the hydrophobic interface to FFA engagement with the receptor, we designed four phenylalanine substitutions in TAS2R14, namely, an F76Y^ECL1^, F76A^ECL1^, F82A^3.25^ and F172A^ECL2^, and tested the ability of the receptor to stimulate the G-protein complex in the presence of FFA through our BRET signaling assay (**Fig. 2E-F**). Our results indicate that these point mutations did not significantly alter the receptor’s activation, either because a single alanine substitution did not have a major effect on the overall hydrophobic nature of the binding pocket, or because the second binding pocket compensated for these changes. The secondary pocket will be discussed extensively in the next section. Notably, alanine substitution at position F82^3.25^ did not have a significant effect on signaling. This is unusual, as the 3.25 position in class A GPCRs is known to be a major determinant of the orthosteric binding pocket geometry and is crucial for agonist binding and signal instigation (*24, 35, 36*). While in most GPCRs 3.25 encompasses a cysteine residue engaged in a disulfide bridge, position 3.25 in TAS2Rs is highly variable, suggesting that its function is not conserved in the taste receptor family.

**Fig. 2.**
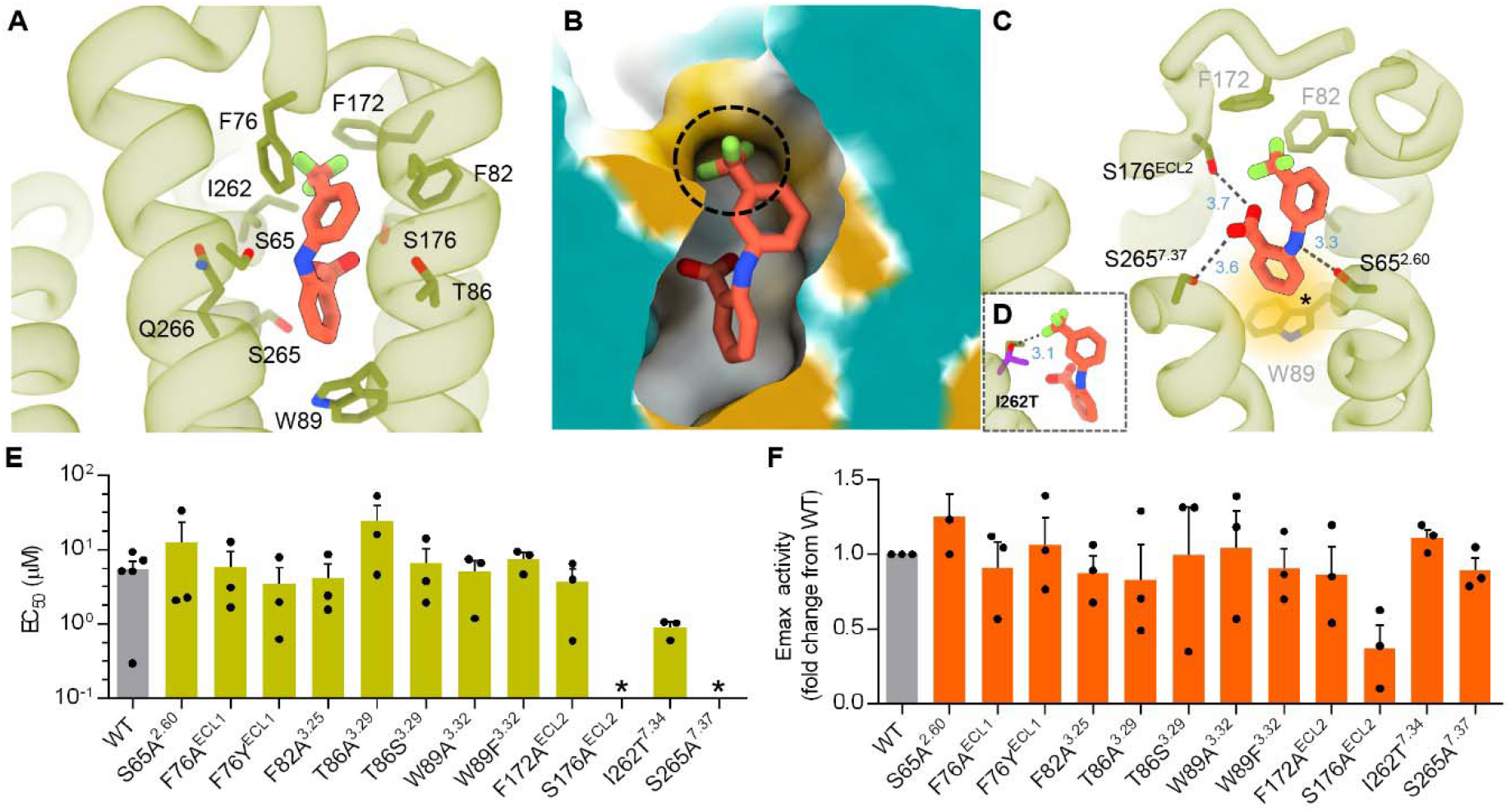
FFA binding at the canonical pocket. (**A**) FFA in the TAS2R14 binding pocket. The receptor is presented in light green, FFA in orange. Residues surrounding the binding pocket are labeled and shown as sticks. (**B**) Ring B of FFA engages a hydrophobic cavity within the canonical binding pocket. Protein surface is mapped by lipophilicity, with hydrophilic residues colored in cyan and lipophilic areas in gold. The area near the trifluoromethyl moiety is circled with a dashed black line. (**C**) FFA interactions in the TAS2R14 binding pocket. Colors are as in A. Dashed lines indicate bonds with distances in Å noted in blue. Perpendicular stacking interactions between ring B and W^3.32^89 are highlighted in yellow and marked by an asterisk. (**D**) An I262T substitution of TAS2R14 contributes to hydrogen bond formation between the fluorine moiety in FFA and the hydroxyl group in threonine and is thus providing a gain of function effect on signaling by FFA. Mutated residue is in purple, the wild-type isoleucine in green. The calculated bond length is indicated in Å. (**E**) EC_50_ values in response to mutations in the TAS2R14 canonical binding pocket. Mutations marked with * in EC50 summary data were not applicable to curve fitting. (**F**) Maximum efficacy (Emax) in fold change from wild-type. Data are represented as mean ±SEM from minimum three independent experiments performed in triplicate.

To further confirm the ligand’s binding pose within the pocket, we designed a gain of function mutation by substituting I262^7.34^ with threonine (I262T^7.34^), to allow for hydrogen bond formation between the hydroxy group in threonine and the fluorine moiety (**Fig. 2D**). Indeed, the BRET assay illustrates a significant decrease in the EC_50_ value of the mutant, suggesting an impact on affinity or on the ability to trigger G-protein activation which displays as a gain of function mutation (**Fig. 2E**).

Ring A in FFA harbors a carboxylic acid moiety that is facing a cavity enriched with polar residues, including S65^2.60^, T86^3.29^, S176^ECL2^, S265^7.38^ and Q266^7.39^, and maintains direct contacts with the serine residues 176 and 265 of ECL2 and TM 7, respectively (**Fig. 2C**). S65^2.60^ makes polar interactions with the amine linker that connects A and B rings in FFA (**Fig. 2C**).

To test the influence of polar contacts on FFA binding, we designed a set of mutations, namely, S65A^2.60^, S176A^ECL2^ and S265A^7.37^. Additionally, we added T86^3.29^A/S mutations, that in previous studies demonstrated significantly reduced susceptibility to FFA (*37*). In our assay, mutation of T86^3.29^ to alanine had a modest effect on Emax but significantly elevated EC_50_, while serine substitution at the same position, did not affect the receptor’s response to FFA (**Fig. 2E-F**). This suggests that despite not having a direct contact with the ligand in the pose observed by cryo-EM, the polar nature of this position is important for FFA activation of TAS2R14. An S176^ECL2^ substitution to alanine reduced the Emax and although the curve fit would not allow an accurate calculation of EC_50_, the data supports an increase in EC_50_ to indicate a significant loss of function (**Fig. 2E-F**). The S265A^7.37^ mutation maintains activity but has a significant elevation in the EC_50_, suggesting this hydrogen bond is crucial for ligand binding (**Fig. 2E-F**). In contrast, an S65A^2.60^ mutation did not gravely impact activation by FFA when compared with the wild-type receptor, implying that the interactions with the amino linker are less essential for ligand binding (**Fig. 2E-F**).

Ring A is further extended towards the receptor’s core and maintains perpendicular aromatic stacking with W89^3.32^ (**Fig. 2C**). W^3.32^ is mostly conserved in TAS2Rs (84%) and is thought to play an important role in receptor activation (*21*). A previous work by Nowak *et al*. (*37*) showed limited effect of the TAS2R14 W89A^3.32^ mutation on the EC_50_ of FFA while an W89F^3.32^ substitution remained unaffected. Similarly, our BRET assays do not show a significant change in response to FFA in the 3.32 mutants (**Fig. 2E-F**). In addition, no change in receptor expression levels was observed while compared with the wild-type, demonstrating that despite the high level of conservation, substitutions at position 3.32 in TAS2R14 did not alter the receptor’s ability to fold, migrate to the cell surface or signal. Notably, an alanine substitution at the synonymous positions in TAS2R10, TAS2R38, TAS2R44 and TAS2R46 impaired the receptors’ responses to a large array of tastants (*21, 38, 39*). This data suggests that compared with other members of the TAS2R family, W^3.32^ in TAS2R14 does not seem to play a significant role in receptor activation by at least some of its ligands, such as FFA. A plausible explanation for the discrepancy could be the second binding pocket for FFA, found through these studies, that could serve as an alternative mechanism for receptor activation.

### Description of FFA binding to the intracellular binding site

In addition to the ligand at the canonical site, we found an unassigned density that bridged the αN5 of gustducin with TMs 3,5 and 7 (**Figs. 1E, 3**). Surprisingly, a copy of FFA could be fitted unambiguously to the density while occupying a conformation that is distinct to the canonical pocket (**Fig. 1E,F**). In this conformation, ring A in the intracellular location is rotated 64° with respect to the same ring at the canonical site (**Fig. 1F**). This binding pose was further validated by Gemspot (*40*) and was also found to be stable through our MD simulations, when compared with alternative poses. In this site, the trifluoromethyl moiety is oriented towards the intracellular face of the receptor and is surrounded with hydrophobic residues of TMs 3, 5 and 7, namely: I111^3.54^, F198^5.58^, L201^5.61^, I202^5.62^, V230^6.35^ and V233^6.38^ (**Fig. 3A-B**). This hydrophobic cavity is further capped with L353^gus-5.25^ of Gα gustducin (residue numbering in gustducin is corresponding to the wild-type human gustducin sequence), and with Y107^3.50^. The carboxyl moiety in ring A is maintaining hydrogen bonds with S194^5.54^ and H276^7.49^.

**Fig. 3.**
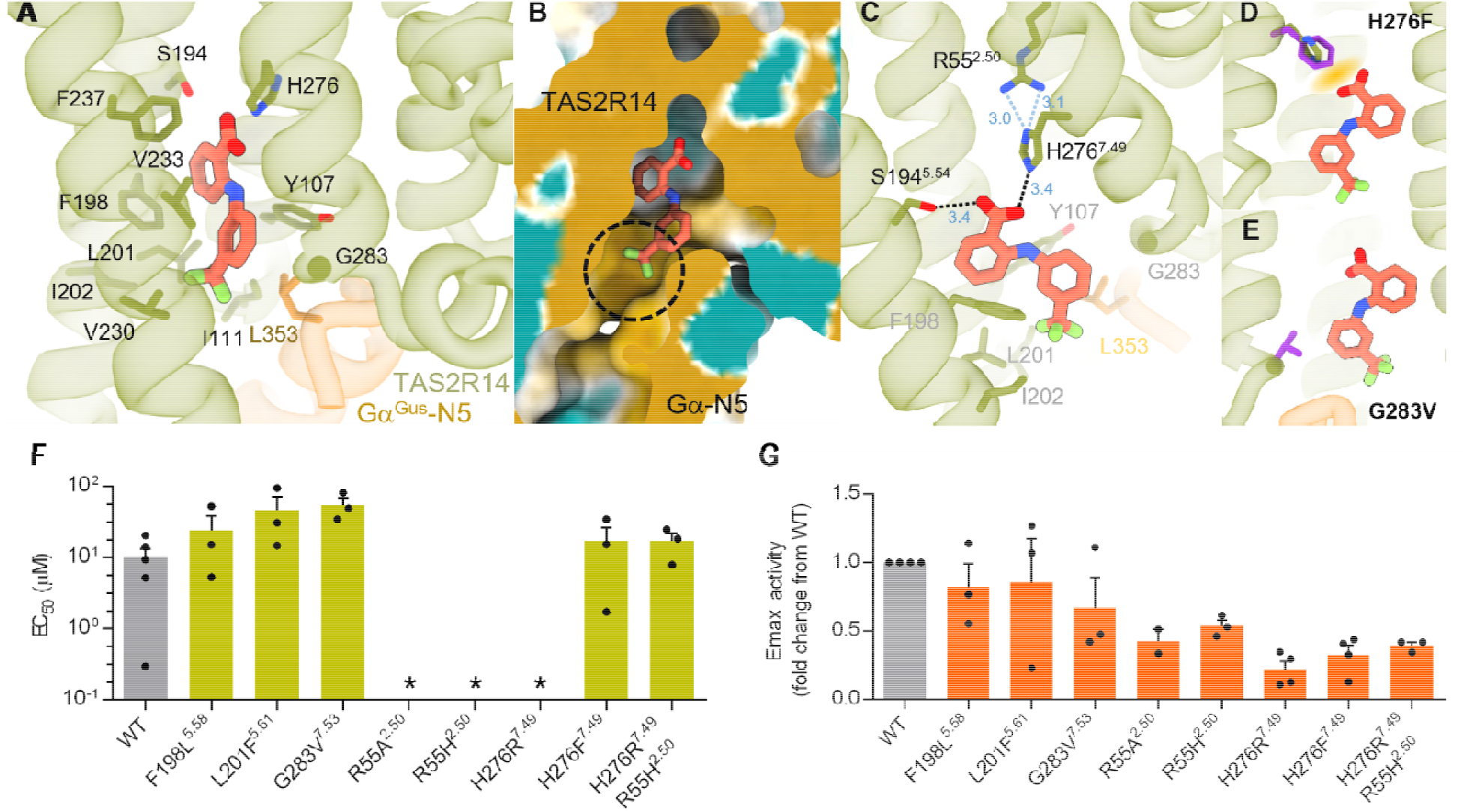
FFA binding in the intracellular pocket. (**A**) FFA binding pose at the intracellular site. The receptor is colored light green, gustducin in gold and FFA in orange. Residues surrounding the binding pocket are labeled and shown as sticks. (**B**) Ring B of FFA engages a hydrophobic pocket composed of TMs 3, 5 and 7 and the α5 helix of gustducin. Protein surface is mapped by lipophilicity, with hydrophilic residues in cyan and lipophilic areas in gold. The α5 helix of gustducin is labeled. The hydrophobic pocket surrounding the trifluoromethyl moiety in FFA is circled. (**C**) FFA interactions in the TAS2R14 binding pocket. Protein and FFA colors are as in panel A. Dashed lines indicate hydrogen bonds with distances in Å noted in blue. (**D**) An H276F mutation is substituting the hydrogen bond between the carboxyl moiety in FFA and the amino group histidine with perpendicular aromatic COOH interactions, and thus only results in partial loss of function. (**E**) A G283V substitution is sterically restricting FFA binding in the intracellular binding pocket. (**F**) EC_50_ values in response to mutations in the TAS2R14 intracellular binding pocket. Mutations marked with an asterisk were not applicable to curve fitting. (**F**) Maximum efficacy (Emax) in fold change from wild-type. Data are represented as mean ±SEM from minimum three independent experiments performed in triplicate

We selected a set of five mutations in residues composing the secondary binding pocket in TAS2R14, F198L^5.58^, L201F^5.61^, G283V^7.53^, H276F^7.49^ and H276R^7.49^ (**Fig. 3F-G**), and performed BRET assays to test the receptor susceptibility to FFA. The first two mutations, F198L^5.58^ (rs202123922) and L201F^5.61^ (rs35804287) are natural variants in human, where L201F^5.61^ was reported to reduce the receptor’s sensitivity to agonists (*13*). The corresponding positions in TAS2Rs are mostly conserved. In our studies, we observed a modest effect on receptor activity with an increase in EC_50_ values for both mutations (**Fig. 3F-G**). In contrast, mutating H276^7.49^ to arginine resulted in complete loss of function, whereas a phenylalanine substitution had significantly lowered Emax with EC_50_ values that were similar to the wild-type receptor activity. The partial activity maintained by phenylalanine could be explained by compensating aromatic-COOH interactions between the carboxyl moiety in ring A and the aromatic ring of phenylalanine (*41*). Notably, position 7.49 is a conserved histidine residue in 24 out of 25 TAS2Rs and composes the H^7.49^S^7.50^xxL^7.53^ motif, that is synonymous to N^7.49^P^7.50^xxY^7.53^ in class A GPCRs (*42*). Our structure further showed that H^7.49^ directly contacts R55^2.50^ (**Fig. 3C**), also conserved in 24 TAS2Rs. 2.50 position in class A receptors is a conserved acidic (mostly D^2.50^) residue known to mediate interactions with sodium ions and to serve as a microswitch for receptor activation (*43, 44*). The high conservation level at these positions in TAS2Rs and their localization within the receptor, suggested that the 2.50-7.49 axis in TAS2Rs can serve as a surrogate microswitch for receptor activity, and thus the reduced sensitivity to FFA observed in our mutational studies might indicate the partial- or full-loss of function due to interference with a key microswitch rather than through ligand binding. To further explore this hypothesis, we designed single and double mutations, an R55A^5.50^, R55H^2.50^ and R55H^2.50^-H276R^7.49^ substitutions. Similar to the H276R^7.49^ mutant, the two single mutations at position 55^2.50^ showed a complete loss of function (**Fig. 3F-G**), supporting the notion that abrogation of the contact with position 276^7.49^ is detrimental to activation by FFA. However, the double mutant, designed to mimic the 2.50-7.49 interaction, was able to partially recover receptor activity with reduced Emax values but an EC_50_ that is similar to the wild-type receptor (**Fig. 3F-G**). These data suggest a key role for the 2.50-7.49 axis. The reduced efficacy in the double mutant could result from the loss of a critical bond with FFA in the intracellular binding pocket, while the canonical site is still intact.

To further explore ligand binding to the intracellular cavity, we designed a loss of function mutant, in a non-conserved residue G283^7.53^. Namely, a G283V^7.53^ was created to in an attempt to restrict the binding pocket to preclude FFA binding (**Fig. 3E**). In this case, we observed a reduction in activation as well as a significant increase in EC_50_ (**Fig. 3F-G**). This result is consistent with our structural observations and further supports the importance of binding of FFA to the intracellular site.

To directly investigate the binding of FFA to TAS2R14, a BRET-based ligand binding assay (*45*) was established. In these experiments we fused TAS2R14 to the small and bright nanoluciferase (Nluc) enzyme (*46, 47*), either at the receptor’s N- or C-terminus, and designed a set of fluorescent FFA derivatives to report proximity to either one of the binding pockets (**Fig. 4A-C**). Our initial experiments were set for the identification of a suitable fluorescent FFA derivative. Based on the available structure-activity relationships (SARs) (*22, 48*), we designed the installation of the fluorophore to be via the meta- or para-positions of the aniline ring B of FFA. Hence, we synthesized a small set of fluorescent derivatives, that comprised linkers of different lengths and polarities that were connected to an azide- or alkyne-functionalized tetramethyl-rhodamine dye (TAMRA-5 or TAMRA-6) employing click chemistry (*49*). This fluorophore has been successfully employed for NanoBRET-based ligand binding assays at intra- and extracellular binding pockets of various GPCRs (*45, 50–52*). We then screened membranes from HEK293T cells that were over-expressing the N- or C-terminally labeled TAS2R14-nanoluciferase constructs and identified TP46 as the most promising candidate specifically binding to both, the intra- and extracellular binding pockets. Saturation binding experiments with TP46 and the extracellularly labelled Nluc-TAS2R14 or the intracellularly labelled TAS2R14-Nluc construct showed typical hyperbolic ligand binding profiles and revealed similar affinities of TP46 for both receptor-Nluc fusion proteins (*K*_D_^extracellular^ 1.80μM; *K*_D_^intracellular^ 1.87μM; **Fig. 4D,E**). Employing TP46 as the molecular probe in competition binding experiments, we determined the affinity of FFA to both ligand binding sites of TAS2R14. Displacement of TP46 with FFA was detected for both TAS2R14-Nluc fusion receptors, demonstrating binding affinities in the low micromolar range of FFA for the both the extracellular (*K*_i_ 11 μM) and the intracellular (*K*_i_ 8.3 μM) sites (**Fig. 4F,G**). The experiments show that the TAMRA-labeled probe TP46 is capable of binding to two distinct sites of TAS2R14 which was detected by nanoBRET with N-terminal and C-terminal receptor-nanoluciferase fusion proteins, respectively. Our competition assays further showed that FFA specifically displaces TP46 and results in a reduced BRET signal from both TAS2R14-NLuc constructs, and thus directly support our observations that FFA binds TAS2R14 at both the canonical and the intracellular pockets.

**Fig. 4.**
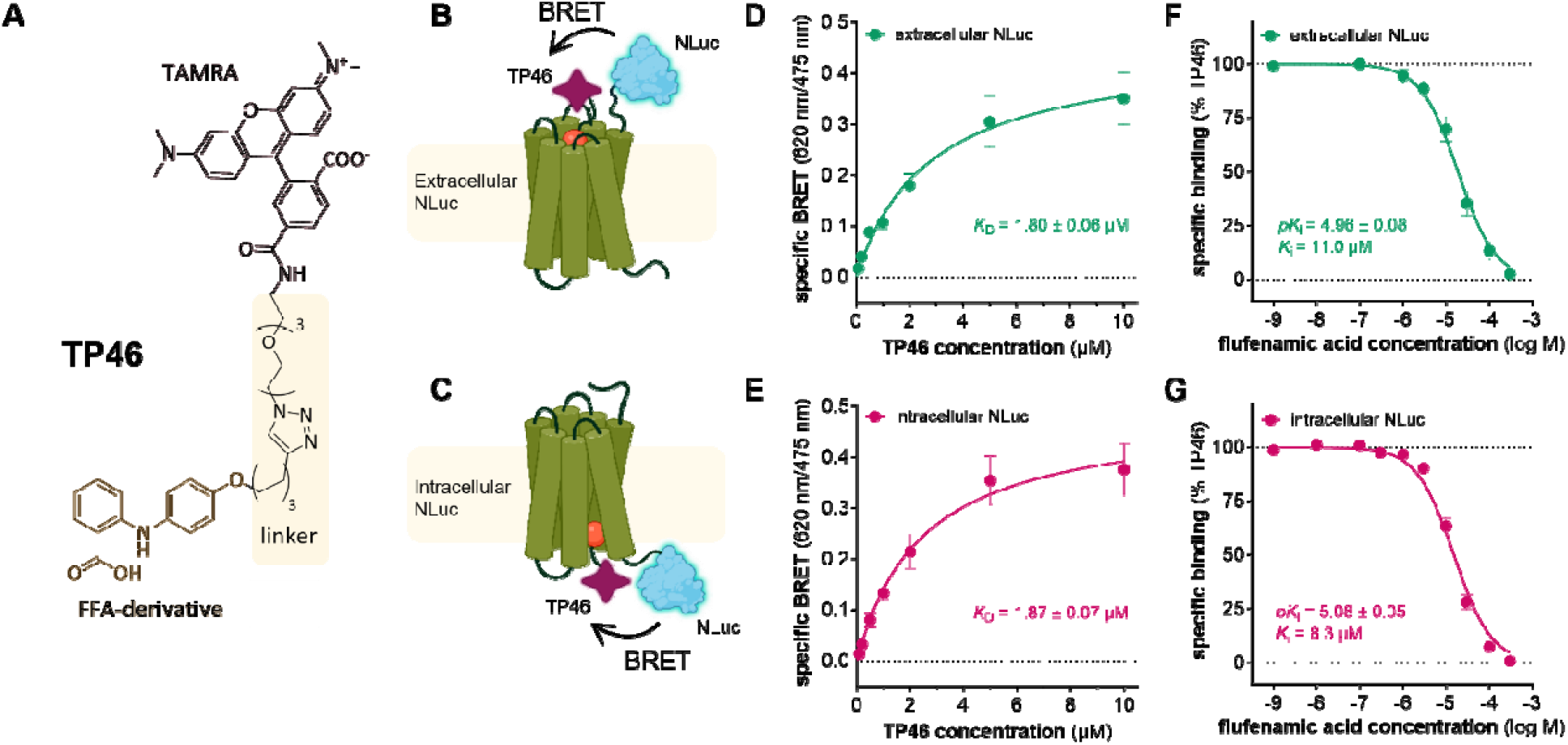
NanoBRET-based ligand binding assays for the extra- and intracellular binding pockets of TAS2R14. (**A**) Chemical structure of the fluorescent molecular probe TP46. (**B-C**) Schematic representation of the NanoBRET-based assay principle for the (**B**) extracellular and (**C**) intracellular NLuc-labeled TAS2R14. (**D-E**) Specific binding of TP46 determined in NanoBRET saturation binding assays reveals similar affinities for the intra- and extracellular binding pockets. Specific BRET was calculated as the difference in total BRET and non-specific binding determined in the presence of 30 μM FFA. Data are shown as mean ± SEM derived from n = 5 (extracellular, **D**) or n = 7 (intracellular, **E**) individual experiments. (**F-G**) Competition binding experiments with TP46 and FFA show binding affinities (K_i_) of 11 μM and 8.3 μM, for the extracellular (**F**) and intracellular (**E**) sites, respectively. Data were normalized to total (100 %) and non-specific BRET (0%) and show mean ± SEM of n = 7 and 16 individual experiments, for the extracellular and intracellular NLuc constructs, respectively.

### Comparison of TAS2R14 with other taste receptors

TAS2Rs are a highly diverse and promiscuous family of receptors that responds to a large number of variable tastants (*4*). A recent study determined the structure of another bitter taste receptor, TAS2R46, in complex with the alkaloid, strychnine (*21*). The comparison of the two structures shows that despite belonging to the same sub-family, the receptors share very little in regards to ligand binding (**Fig. 5**).

**Fig. 5.**
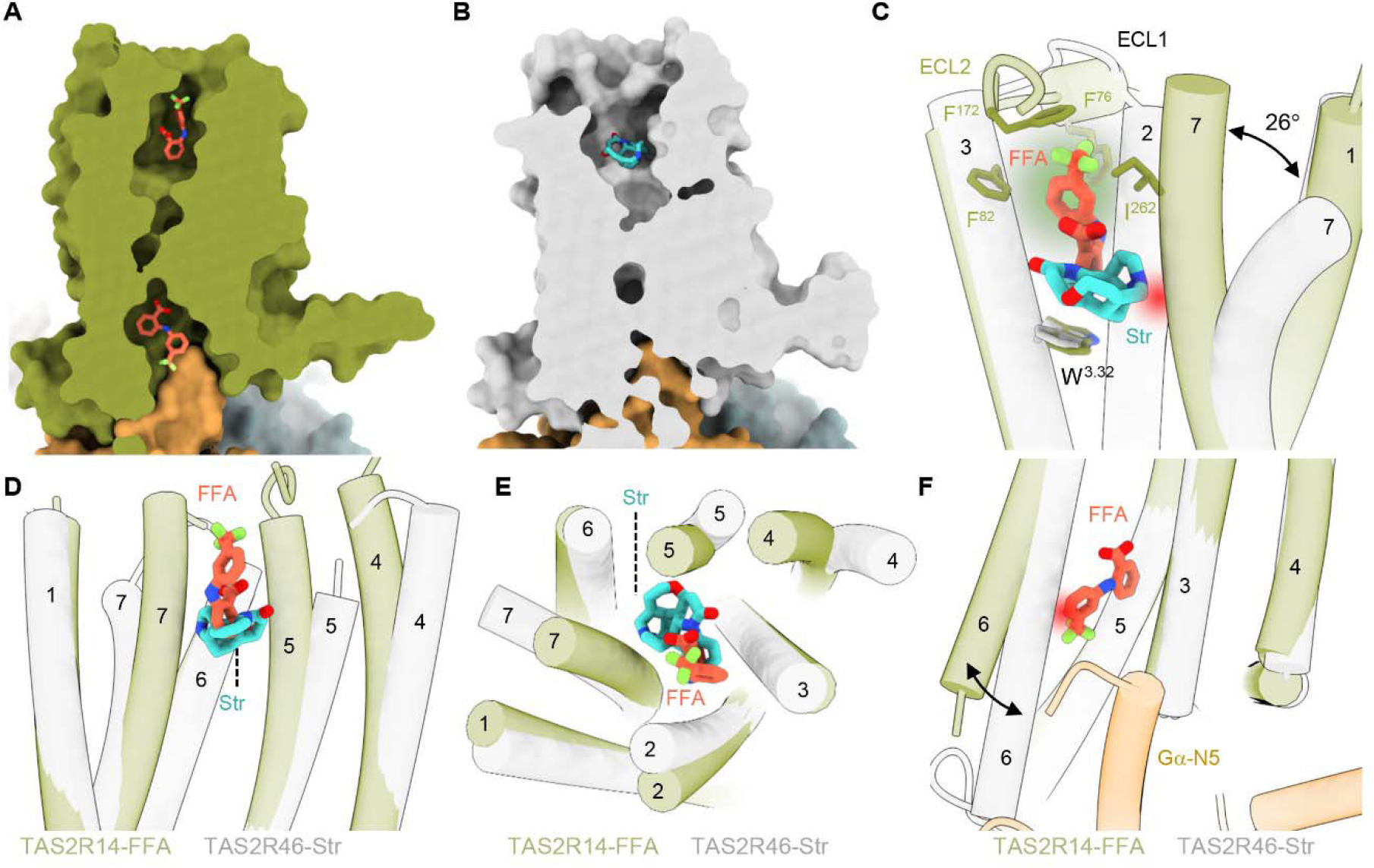
Comparison of TAS2R14 with TAS2R46. (**A-B**) A view of FFA (orange, **A**) and strychnine (Str, cyan, PDB 7XP6, **B**) at the TA2R binding pocket. TAS2R14 is in green (**A**), and TAS2R46 in gray (**B**) Gustducin is shown in gold for both receptors. (**C**) Superposition of FFA and Str within the canonical binding pocket reveals different binding modes for the two ligands. FFA extends into a cavity localized towards the extracellular milieu. The cavity is highlighted in green and composes the hydrophobic pocket that accommodates the trifluoromethyl moiety. Str clashes with the inward TM7 position of TAS2R14, thus there is an outward movement of TM7 at the extracellular site. Clash is presented in red. Residue and helix numbers are indicated in the figure. TAS2R14 is green, TAS2R46 is grey. (**D-E**) Comparison of TM arrangement in the canonical binding pocket reveals major changes in the orientation of TMs 2,4,5 and 7 in the bound structures of TAS2R14 (green) and TAS2R46 (gray). A side view is in (**D**), top view is presented in (**E**). (**F**) Superposition of TAS2R14 (green) and TAS2R46 (gray) reveals that in the intracellular face, TM arrangement is identical for both receptors, apart from an outward movement of TM6 in TAS2R14. A clash with TM6 position in TAS2R46 (that lacks the second pocket) is indicated in red.

At the canonical pocket, the structures greatly differ both in shape and volume of the cavity; While TAS2R14 has a rather narrow void, TAS2R46 has a wide opening that is also more exposed to the extracellular milieu (surface areas are 511- and 426-Å^3^ for TAS2R46 and TAS2R14, respectively; **Fig. 5A-B**). Within the pocket, FFA and strychnine, that differ in their chemical structure and composition, have limited overlap and occupy different regions of their respected cavities (**Fig. 5C**). Strychnine has an overall spherical shape and is packed at the bottom of the pocket stacking against a conserved tryptophan residue at position 3.32 (W88^3.32^ in TAS2R46). In contrast, FFA, that has an elongated shape, is oriented in a perpendicular orientation (maintaining interactions with the corresponding residue, W89^3.32^, in TAS2R14) and extends towards the extracellular space while interacting with a hydrophobic cavity composed of non-conserved residues of ECL1, ECL2 and the tips of TMs 3 and 7 (**Fig. 5C**). This unique cavity is formed by different arrangement of the extracellular surroundings in TAS2R14, that includes the packing of the extracellular loops towards the receptor’s core along with an inward-inclined arrangement of TMs 4, 5 and 7 at the extracellular face (**Fig. 5C-E**). TM2 is also slightly shifted outwards in comparison with TAS2R46 structure, creating space that accommodates ring B of FFA (**Fig. 5E**).

The most notable difference in the extracellular TM arrangement is the TM7 position (**Figs. 5C**). In both receptors TM7 has a kink, that is caused by a hinge originating from a shared proline residue in position 7.46 (P273^7.46^ and P272^7.46^ in TAS2R14 and TAS2R46, respectively). However, the tilt level greatly differs, with only a mild tilt (22°) for TAS2R14, and a 45° tilt in TAS2R46, that profoundly opens the extracellular space of the receptor and contributes to the larger void of the binding pocket (**Figs 5C-E** ). Considering the differential binding mode of the two ligands within the pockets, it is tempting to speculate that the tilt level might depend on the ligand’s geometry, as strychnine, that extends towards TM7, clashes with the TM7 position in TAS2R14. FFA, on the other hand, extends upwards and maintains interactions with I262^7.34^ that is located at the tip of TM7, and may stabilize the orientation of TM7 closer to the 7TM core (**Fig. 5C**). Also, given that the proline residue (P^7.46^), that serves as a hinge to enable TM7 mobilization, is conserved in most TAS2Rs, but not in other GPCRs, this could represent a common mechanism for bitter taste receptors. In fact, the odorant receptor family is the only other GPCR family with conserved P^7.46^. This family of receptors also recognizes numerous and diverse ligands, which could further support the role of P^7.46^ for versatility. In addition, despite the large displacement of TM7 at the extracellular facet, the intracellular part of TM7 does not differ between the two receptors. This would further suggest that the proline hinge can dictate a unidirectional movement of the helix to allow the accommodation of ligands with different diameter at the extracellular site, while keeping the intracellular site at a conformation that will allow the interaction with the intracellular partners.

The most profound difference between the two receptors is that in TAS2R46, the ligand only binds at the extracellular pocket, and no ligand is present in the intracellular site (**Fig. 5A-B**). Comparing the intracellular domain of the two receptors at the structural level, it is noticeable that while the extracellular sites exhibit a pronouncedly different TM rearrangement, the intracellular face is nearly identical, apart from an apparent displacement of TM6 outwards in the TAS2R14 structure (**Fig. 5F**). Notably, albeit the residues that are in direct contact with the FFA molecule at the intracellular site are conserved, one significant difference is that position 7.50 in TAS2R14 is a serine (S277^7.50^), while at TAS2R46, the orthologous residue is proline (P276^7.50^). This position is a conserved proline in most GPCRs and composes the N^7.49^P^7.50^xxY^7.53^ motif, a key signature motif in class A receptors that is crucial for the structural rearrangements of receptors during activation. In TAS2Rs, this motif is replaced by an H^7.49^S/P^7.50^xxL^7.53^ consensus (*42*). Position 7.50 is not in direct contact with FFA, but H^7.49^, also part of the motif, is maintaining polar contacts with the ligand, and our mutational studies confirm the importance of this position for signal transduction (**Fig. 3F-G**). However, the arrangement of the TMs, and in particular of TM6, is much tighter in the currently available structures of TAS2R46, which have no sufficient volume for an intracellular ligand binding pocket. In regards with the G protein contact, the overall conformation of the two receptors is nearly identical around the FFA secondary binding site, and so are the interactions with gustducin. However, at a more global level, the TM6 movement in TAS2R14 breaks two contacts between the receptor and the α5 helix of gustducin. Those include S220^6.27^ in TAS2R46 that interacts with D341^gus-H5.13^ and K345^gus-H5.17^ and V223^6.30^ that interacts with F354^gus-H5.26^. While these positions are mostly conserved in TAS2Rs, they correspond to A222^6.27^ and K225^6.30^ in TAS2R14, respectively. Although these contacts are remote from the binding pocket, the fact that they are different in TAS2R14 compared with other TAS2Rs might change the tendency of TAS2R14 to bind gustducin at the interface, rendering TM6 more mobile and thus possibly more prone for ligand entry to the secondary site.

This analysis shows that despite belonging to the same receptor class, the binding mode of tastants to bitter taste receptors is highly diversified. A conservation analysis comparing all human class T receptors similarly illustrates that the extracellular pocket of TAS2R is highly variable amongst human TAS2Rs with the exception of W^3.32^, while the intracellular site is more conserved. This could be explained by the need to have a rather diverse extracellular pocket for sensing a large array of ligands with varying chemical scaffolds, along with a joint mechanism with the intracellular partners. The potential for at least some of the TAS2Rs to accommodate ligands at the intracellular site adds to the versatility and complexity of the TAS2R family.

### Comparing the intracellular pocket with other reported intracellular ligands

Recent studies revealed the binding of small molecular ligands at the intracellular face of other GPCRs. Those include several class A members, the chemokine receptors (CCRs 2,7 and 9 (*53–55*)), CXCR2 (*56*), the orphan receptor GPR61 (*57*), the β2 adrenergic receptor (*58*), and the neurotensin 1 receptor (NTSR1) (*59*), as well as the parathyroid 1 (PTH1R) (*60*) and the corticotropin releasing factor type 1 (CRF1R) (*61*) receptors, of class B. In addition, a palmitoyl moiety that is anchored to the αN5 helix of Gα_o_ was also reported to occupy the same cavity in a structure of the adhesion GPCR (class B2), GPR97 (*62*) .

Superposition of the bound structures reveals that despite bearing very different chemical scaffolds, the ligands’ positions largely overlap, where they jointly engage residues in TMs 3,5,6 and 7 of their respective receptors (**Fig. 6A-B**). Within this relatively small set of compounds, class-A binders extend towards the G protein-receptor interface, thus engaging with unique positions in TM1 and H8; whereas class B ligands are pointing towards the canonical binding site, maintaining interactions with residues at TMs 5 and 6 that are deeply penetrating the 7TM core towards the canonical binding site (**Fig. 6A-B**). As a result, the class A binders interact with both the D[E]^3.49^R^3.50^Y^3.51^ and the N^7.49^P^7.50^xxY^7.53^ motifs via R^3.50^ and Y^7.53^, respectively (**Fig. 6C**); these motifs are missing in class B. In contrast, class B ligands uniquely target the P^6.47b^xxG^6.50b^ motif that was found to be key for the structural rearrangement required for the activation of this class of receptors (**Fig. 6C**). FFA interactions with the TAS2R14 binding pocket mostly resemble class A ligands (**Fig. 6C-D**). Notable common features are the interaction with Y^3.50^ and H^7.49^ of the F^3.49^Y^3.50^xxK^3.53^ and the H^7.49^S/P^7.50^xxL^7.53^ motifs, that are analogous to the D[E]^3.49^R^3.50^Y^3.51^ and the N^7.49^P^7.50^xxY^7.53^ motifs in class A, respectively (*42*). A main difference between TAS2R14 and the other reported structures is that in the other structures, the ligands were only bound at the intracellular site, and no ligands were observed at the canonical binding pocket, except for the GPR97 structure, where in addition to the palmitoyl moiety, occupying the intracellular cavity, there is a sterol partial agonist bound at the canonical pocket (*62*). Nevertheless, the physiological significance of this binding is still unclear.

**Fig. 6.**
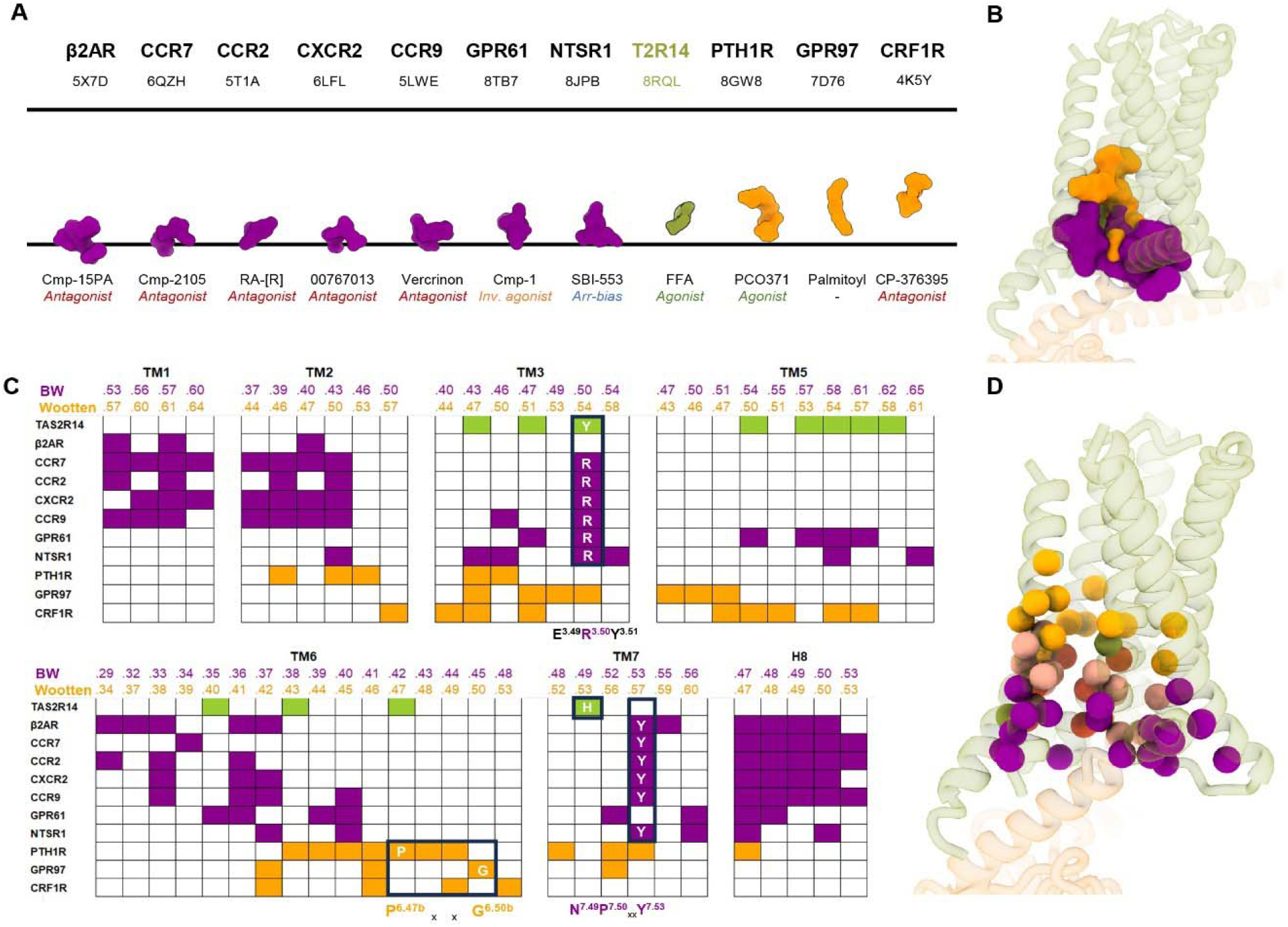
Comparison of the TAS2R secondary binding site with intracellular pockets of other GPCRs. (**A**) Localization of ligands targeting intracellular binding pockets in GPCRs. The black lines represent the membrane boundaries and are used as a reference to show that different ligands occupy slightly different interfaces within the intracellular cavity. Colors indicate family affiliation, with class A in magenta, class T (taste) in green, and class B1,B2 in gold. GPCR identity and PDB ID are labeled on top of the scheme. The bottom labels indicate the ligand name and reported activity (agonist/antagonist/inverse agonist). (**B**) Representation of ligand position in the intracellular site in 3D. Structures were superposed to TAS2R14 (aligned on TM4). Similar to the 2D scheme in (A), the superposition shows that class B ligands (gold) occupy a higher position in the pocket when compared with class A binders (magenta). FFA (green) is similar to class A in regard to the positioning within the pocket. (**C**) Characterization of the residues within the intracellular binding pocket. Residues shown are within 4 Å distance from their respective ligands. Receptor names are indicated to the left; BW (class A) and Wootten (class B) numbering of residues in their respective TMs are on top of each column. Colors are according to family affiliation and are as in A. Conserved positions or positions that appear in most pockets are also indicated by the relevant amino-acid letter code. (**D**) Representation of residues involved in ligand binding in the intracellular pocket in 3D. Residues that are strictly interacting with class A, B or TAS2R14 are in their respected colors (magenta, gold and green). Residues that participate in binding of all reported ligands are in red. Residues that participate in more than one receptor class, but not all, are in pink.

In these regards, FFA seems to mimic the intracellular binding of class A agonists. Altogether, the intracellular pocket of GPCRs might be a widespread feature that so far was reported for only a few members of class A, B and in this work. This could expand the possibilities of current hotspots in GPCR targeting, and may be of an immense pharmacological interest.

## Summary

Our cryo-EM structure of TAS2R14 reveals two binding modes for FFA, one at the canonical binding site and another at the intracellular facet, where it interacts with key elements important for GPCR activation and with the intracellular partner gustducin. Our novel binding assay shows that FFA binds at both sites with similar affinities that are at the micromolar range. Functional studies of the intracellular binding site also establish a key role for the H276^7.49^ and R55^2.50^ positions in receptor activation. Due to their conservation in the TAS2R family, they might serve as a family-specific activation axis. The intracellular binding pocket, similar to the pocket recently reported for class A and B GPCRs is highlighted as a hotspot for GPCR targeting, that may further expand the pharmacological space for targeted drug design. Our work provides the tools for site-directed structure guided design for bitter taste GPCRs, which are not only key players for food sensing and evaluation but are also expressed extra-orally and have roles in several diseases and disorders.

## Acknowledgments

We thank Khajidmaa Flad for the cloning of the secNLuc-TAS2R14 construct, Asa Tirosh, Evgenii Ziaikin and George Chalhoub for the helpful discussions. We thank Nadav Elad for technical support with microscope operation.

## References

1. W. Meyerhof, C. Batram, C. Kuhn, A. Brockhoff, E. Chudoba, B. Bufe, G. Appendino, M. Behrens, The molecular receptive ranges of human TAS2R bitter taste receptors. Chem. Senses 35, 157–170 (2009).

2. K. Lossow, S. Hübner, N. Roudnitzky, J. P. Slack, F. Pollastro, M. Behrens, W. Meyerhof, Comprehensive Analysis of Mouse Bitter Taste Receptors Reveals Different Molecular Receptive Ranges for Orthologous Receptors in Mice and Humans. J. Biol. Chem. 291, 15358–15377 (2016).

3. A. Di Pizio, M. Y. Niv, Promiscuity and selectivity of bitter molecules and their receptors. Bioorganic Med. Chem. 23, 4082–4091 (2015).

4. A. Dagan-wiener, A. Di Pizio, I. Nissim, M. S. Bahia, N. Dubovski, E. Margulis, M. Y. Niv, BitterDB□: taste ligands and receptors database in 2019. 47, 1179–1185 (2019).

5. T. Delompré, C. Belloir, C. Martin, C. Salles, L. Briand, Detection of Bitterness in Vitamins Is Mediated by the Activation of Bitter Taste Receptors. Nutrients 14 (2022).

6. W. S. U. Roland, L. Van Buren, H. Gruppen, M. Driesse, R. J. Gouka, G. Smit, J. P. Vincken, Bitter taste receptor activation by flavonoids and isoflavonoids: Modeled structural requirements for activation of hTAS2R14 and hTAS2R39. J. Agric. Food Chem. 61, 10454–10466 (2013).

7. M. Behrens, A. Brockhoff, C. Kuhn, B. Bufe, M. Winnig, W. Meyerhof, The human taste receptor hTAS2R14 responds to a variety of different bitter compounds. Biochem. Biophys. Res. Commun. 319, 479–485 (2004).

8. A. Levit, S. Nowak, M. Peters, A. Wiener, W. Meyerhof, M. Behrens, M. Y. Niv, The bitter pill: Clinical drugs that activate the human bitter taste receptor TAS2R14. FASEB J. 28, 1181–1197 (2014).

9. E. Margulis, T. Lang, A. Tromelin, E. Ziaikin, M. Behrens, M. Y. Niv, Bitter Odorants and Odorous Bitters: Toxicity and Human TAS2R Targets. J. Agric. Food Chem. 71, 9051–9061 (2023).

10. S. Grassin-delyle, C. Abrial, S. Fayad-kobeissi, M. Brollo, C. Faisy, J. Alvarez, E. Naline, P. Devillier, The expression and relaxant effect of bitter taste receptors in human bronchi. 1–14 (2013).

11. C. H. Yan, S. Hahn, D. Mcmahon, D. Ph, D. Bonislawski, D. W. Kennedy, N. D. Adappa, J. N. Palmer, P. Jiang, D. Ph, R. J. Lee, D. Ph, N. A. Cohen, D. Ph, Nitric oxide production is stimulated by bitter taste receptors ubiquitously expressed in the sinonasal cavity. Am. J. Rhinol. Allergy 38, 85–92 (2017).

12. C. J. Bloxham, S. R. Foster, W. G. Thomas, W. G. Thomas, A Bitter Taste in Your Heart. Front. Physiol. 11 (2020).

13. C. J. Bloxham, K. D. Hulme, F. Fierro, C. Fercher, C. L. Pegg, S. L. O. Brien, S. R. Foster, K. R. Short, S. G. B. Furness, M. E. Reichelt, M. Y. Niv, G. Walter, Cardiac human bitter taste receptors contain naturally occurring variants that alter function. Biochem. Pharmacol., 115932 (2023).

14. K. I. Liszt, J. P. Ley, B. Lieder, M. Behrens, V. Stöger, A. Reiner, C. M. Hochkogler, E. Köck, A. Marchiori, J. Hans, S. Widder, G. Krammer, G. J. Sanger, M. M. Somoza, W. Meyerhof, V. Somoza, Caffeine induces gastric acid secretion via bitter taste signaling in gastric parietal cells. Proc. Natl. Acad. Sci. U. S. A. 114, E6260–E6269 (2017).

15. M. Gentiluomo, L. Crifasi, A. Luddi, D. Locci, R. Barale, P. Piomboni, D. Campa, Taste receptor polymorphisms and male infertility. Hum. Reprod. 32, 2324–2331 (2017).

16. A. Jaggupilli, N. Singh, J. Upadhyaya, A. S. Sikarwar, M. Arakawa, S. Dakshinamurti, R. P. Bhullar, K. Duan, P. Chelikani, Analysis of the expression of human bitter taste receptors in extraoral tissues. Mol. Cell. Biochem. 426, 137–147 (2017).

17. N. Singh, F. A. Shaik, Y. Myal, P. Chelikani, Chemosensory bitter taste receptors T2R4 and T2R14 activation attenuates proliferation and migration of breast cancer cells. Mol. Cell. Biochem. 465, 199–214 (2020).

18. Z. A. Miller, J. F. Jolivert, R. Z. Ma, S. Muthuswami, A. Mueller, D. B. Mcmahon, R. M. Carey, R. J. Lee, Lidocaine Induces Apoptosis in Head and Neck Squamous Cell Carcinoma Cells Through Activation of Bitter Taste Receptor T2R14. Cell Rep., 113437 (2023).

19. L. Stern, L. F. Boehme, M. R. Goetz, C. Nitschke, A. Giannou, T. Zhang, C. Güngör, M. Reeh, J. R. Izbicki, R. Fliegert, A. Hausen, N. Giese, T. Hackert, M. Y. Niv, S. Heinrich, M. M. Gaida, T. Ghadban, Potential role of the bitter taste receptor T2R14 in the prolonged survival and enhanced chemoresponsiveness induced by apigenin. Int. J. Oncol. 62, 1–14 (2023).

20. S. K. Mclaughlin, P. J. Mckinnon, R. F. Margolskee, Gustducin is a taste-cell-specific G protein closely related to the transducins. 357, 563–569 (1992).

21. W. Xu, L. Wu, S. Liu, X. Liu, X. Cao, C. Zhou, J. Zhang, Y. Fu, Y. Guo, Y. Wu, Q. Tan, L. Wang, J. Liu, L. Jiang, Z. Fan, Y. Pei, J. Yu, J. Cheng, S. Zhao, X. Hao, Z. Liu, T. Hua, Structural basis for strychnine activation of human bitter taste receptor TAS2R46. Science (80-.). 1303, 1298–1303 (2022).

22. A. Di Pizio, L. A. W. Waterloo, R. Brox, S. Löber, D. Weikert, M. Behrens, P. Gmeiner, M. Y. Niv, Rational design of agonists for bitter taste receptor TAS2R14L: from modeling to bench and back. Cell. Mol. Life Sci., doi: 10.1007/s00018-019-03194-2 (2019).

23. S. Maeda, A. Koehl, H. Matile, H. Hu, D. Hilger, G. F. X. Schertler, A. Manglik, G. Skiniotis, R. J. P. Dawson, B. K. Kobilka, Development of an antibody fragment that stabilizes GPCR/G-protein complexes. Nat. Commun. 9, 1–9 (2018).

24. H. Israeli, O. Degtjarik, F. Fierro, V. Chunilal, A. K. Gill, N. J. Roth, J. Botta, V. Prabahar, Y. Peleg, L. F. Chan, D. Ben-Zvi, P. J. McCormick, M. Y. Niv, M. Shalev-Benami, Structure reveals the activation mechanism of the MC4 receptor to initiate satiation signaling. Science (80-.). 372, 808–814 (2021).

25. K. Kim, T. Che, O. Panova, J. F. DiBerto, J. Lyu, B. E. Krumm, D. Wacker, M. J. Robertson, A. B. Seven, D. E. Nichols, B. K. Shoichet, G. Skiniotis, B. L. Roth, Structure of a Hallucinogen-Activated Gq-Coupled 5-HT2A Serotonin Receptor. Cell 182, 1574–1588.e19 (2020).

26. A. Koehl, H. Hu, S. Maeda, Y. Zhang, Q. Qu, J. M. Paggi, N. R. Latorraca, D. Hilger, R. Dawson, H. Matile, G. F. X. Schertler, S. Granier, W. I. Weis, R. O. Dror, A. Manglik, G. Skiniotis, B. K. Kobilka, Structure of the μ-opioid receptor-Gi protein complex. Nature 558, 547–552 (2018).

27. B. Carpenter, A. Singhal, A. Strege, P. C. Edwards, C. F. White, H. Du, R. Grisshammer, C. G. Tate, Mini-G proteinsL: Novel tools for studying GPCRs in their active conformation. PLoS One 12, 1–26 (2017).

28. A. S. Dixon, M. K. Schwinn, M. P. Hall, K. Zimmerman, P. Otto, T. H. Lubben, B. L. Butler, B. F. Binkowski, T. MacHleidt, T. A. Kirkland, M. G. Wood, C. T. Eggers, L. P. Encell, K. V. Wood, NanoLuc Complementation Reporter Optimized for Accurate Measurement of Protein Interactions in Cells. ACS Chem. Biol. 11, 400–408 (2016).

29. A. Inoue, F. Raimondi, F. M. N. Kadji, G. Singh, T. Kishi, A. Uwamizu, Y. Ono, Y. Shinjo, S. Ishida, N. Arang, K. Kawakami, J. S. Gutkind, J. Aoki, R. B. Russell, Illuminating G-Protein-Coupling Selectivity of GPCRs. Cell 177, 1933–1947.e25 (2019).

30. R. H. J. Olsen, J. F. DiBerto, J. G. English, A. M. Glaudin, B. E. Krumm, S. T. Slocum, T. Che, A. C. Gavin, J. D. McCorvy, B. L. Roth, R. T. Strachan, TRUPATH, an open-source biosensor platform for interrogating the GPCR transducerome. Nat. Chem. Biol. 16, 841–849 (2020).

31. C. Galés, J. J. J. Van Durm, S. Schaak, S. Pontier, Y. Percherancier, M. Audet, H. Paris, M. Bouvier, Probing the activation-promoted structural rearrangements in preassembled receptor – G protein complexes. Nat. Struct. Mol. Biol. 13, 778–786 (2006).

32. J. Duan, D. Shen, X. E. Zhou, P. Bi, Q. Liu, Y. Tan, Y. Zhuang, H. Zhang, P. Xu, S. Huang, S. Ma, X. He, K. Melcher, Y. Zhang, H. E. Xu, Y. Jiang, Cryo-EM structure of an activated VIP1 receptor-G protein complex revealed by a NanoBiT tethering strategy. Nat. Commun. 11, 1–10 (2020).

33. F. Fierro, L. Peri, H. Hübner, A. Tabor-Schkade, L. Waterloo, S. Löber, T. Pfeiffer, D. Weikert, T. Dingjan, E. Margulis, P. Gmeiner, M. Y. Niv, Inhibiting a promiscuous GPCR: iterative discovery of bitter taste receptor ligands. Cell. Mol. Life Sci. 80, 1–17 (2023).

34. J. A. Ballesteros, H. Weinstein, “[19] Integrated methods for the construction of three-dimensional models and computational probing of structure-function relations in G protein-coupled receptors” in Receptor Molecular Biology, S. C. B. T.-M. in N. Sealfon, Ed. (Academic Press, 1995; https://www.sciencedirect.com/science/article/pii/S1043947105800497)vol. 25, xpp. 366–428.

35. K. K. Kumar, M. Shalev-benami, M. J. Robertson, S. V Malhotra, B. K. Kobilka, K. K. Kumar, M. Shalev-benami, M. J. Robertson, H. Hu, S. D. Banister, Structure of a Signaling Cannabinoid Receptor 1-G Article Structure of a Signaling Cannabinoid Receptor 1-G Protein Complex. Cell 176, 448–458.e12 (2019).

36. A. J. Venkatakrishnan, X. Deupi, G. Lebon, C. G. Tate, G. F. Schertler, M. M. Babu, Molecular signatures of G-protein-coupled receptors. Nature 494, 185–194 (2013).

37. S. Nowak, A. Di Pizio, A. Levit, M. Y. Niv, W. Meyerhof, M. Behrens, Reengineering the ligand sensitivity of the broadly tuned human bitter taste receptor TAS2R14. Biochim. Biophys. Acta - Gen. Subj. 1862, 2162–2173 (2018).

38. A. N. Pronin, H. Tang, J. Connor, W. Keung, N. T. P. Road, L. Jolla, Identification of Ligands for Two Human Bitter T2R Receptors. Identif. Ligands Two Hum. Bitter T2R Recept. 29, 583–593 (2004).

39. S. Born, A. Levit, M. Y. Niv, W. Meyerhof, M. Behrens, The Human Bitter Taste Receptor TAS2R10 Is Tailored to Accommodate Numerous Diverse Ligands. J. Neurosci. 33, 201–213 (2013).

40. M. J. Robertson, G. C. P. van Zundert, K. Borrelli, G. Skiniotis, GemSpot: A Pipeline for Robust Modeling of Ligands into Cryo-EM Maps. Structure 28, 707–716.e3 (2020).

41. M. R. Jackson, R. Beahm, S. Duvvuru, C. Narasimhan, J. Wu, H. Wang, V. M. Philip, R. J. Hinde, E. E. Howell, A Preference for Edgewise Interactions between Aromatic Rings and Carboxylate AnionsL: The Biological Relevance of Anion - Quadrupole Interactions. J. Phys. Chem. B 111, 8242–8249 (2007).

42. A. Di Pizio, A. Levitx, M. Slutzki, M. Behrens, R. Karamanjj, M. Y. Niv, Comparing Class A GPCRs to Bitter Taste Receptors: Structural Motifs, Ligand Interactions and Agonist-to-Antagonist Ratios (Elsevier Ltd, 2016; 10.1016/bs.mcb.2015.10.005)vol. 132.

43. S. R, M. R, E. A, N. D, Allosteric sodium in class A GPCR signaling. Trends Biochem Sci 39, 233–244 (2014).

44. D. Hilger, The role of structural dynamics in GPCR-mediated signaling. FEBS J. 288, 2461–2489 (2021).

45. L. A. Stoddart, E. K. M. Johnstone, A. J. Wheal, J. Goulding, M. B. Robers, T. Machleidt, K. V Wood, S. J. Hill, K. D. G. Pfleger, Application of BRET to monitor ligand binding to GPCRs. Nat. Methods 12, 661–663 (2015).

46. M. P. Hall, J. Unch, B. F. Binkowski, M. P. Valley, B. L. Butler, M. G. Wood, P. Otto, K. Zimmerman, G. Vidugiris, T. Machleidt, M. B. Robers, A. Benink, C. T. Eggers, M. R. Slater, P. L. Meisenheimer, D. H. Klaubert, F. Fan, L. P. Encell, K. V Wood, Engineered Luciferase Reporter from a Deep Sea Shrimp Utilizing a Novel Imidazopyrazinone Substrate. Chem. Biol., 1848–1857 (2012).

47. C. G. England, E. B. Ehlerding, W. Cai, NanoLuc: A Small Luciferase Is Brightening Up the Field of Bioluminescence. Bioconjug. Chem. 27, 1175–1187 (2016).

48. L. Waterloo, H. Hu, F. Fierro, T. Pfeiffer, R. Brox, L. Stefan, D. Weikert, M. Y. Niv, P. Gmeiner, Discovery of 2LAminopyrimidines as Potent Agonists for the Bitter Taste Receptor TAS2R14. J. Med. Chem. Fig., doi: 10.1021/acs.jmedchem.2c01997 (2023).

49. V. V Rostovtsev, L. G. Green, V. V Fokin, K. B. Sharpless, A Stepwise Huisgen Cycloaddition ProcessL: Copper (i)-Catalyzed Regioselective TM Ligation ∫ of Azides and Terminal Alkynes **. Angew. Chemie 114, 2596–2599. (2002).

50. M. E. Huber, L. Toy, M. F. Schmidt, H. Vogt, J. Budzinski, M. F. J. Wiefhoff, N. Merten, E. Kostenis, D. Weikert, M. Schiedel, A Chemical Biology Toolbox Targeting the Intracellular Binding Site of CCR9: Fluorescent Ligands, New Drug Leads and PROTACs. Angew. Chemie - Int. Ed. 61, 1–5 (2022).

51. A. Allikalt, N. Purkayastha, K. Flad, M. F. Schmidt, A. Tabor, P. Gmeiner, H. Hübner, D. Weikert, Fluorescent ligands for dopamine D2/D3 receptors. Sci. Rep. 10, 1–13 (2020).

52. N. Rosier, L. Gr, H. Schihada, M. Jan, A. Is, L. J. Humphrys, M. Nagl, U. Seibel, M. J. Lohse, A Versatile Sub-Nanomolar Fluorescent Ligand Enables NanoBRET Binding Studies and Single-Molecule Microscopy at the Histamine H3 Receptor. J. Med. Chem., 11695–11708 (2021).

53. Y. Zheng, L. Qin, V. Natalia, O. Zacarías, H. De Vries, G. W. Han, M. Gustavsson, M. Dabros, C. Zhao, R. J. Cherney, P. Carter, D. Stamos, R. Abagyan, V. Cherezov, R. C. Stevens, Structure of CC chemokine receptor 2 with orthosteric and allosteric antagonists. Nature 540, 458–461 (2016).

54. S. Saha, A. K. Shukla, The Inside Story: Crystal Structure of the Chemokine Receptor CCR7 with an Intracellular Allosteric Antagonist. Biochemistry 59, 12–14 (2020).

55. C. Oswald, M. Rappas, J. Kean, A. S. Doré, J. C. Errey, K. Bennett, F. Deflorian, J. A. Christopher, A. Jazayeri, J. S. Mason, M. Congreve, R. M. Cooke, F. H. Marshall, Intracellular allosteric antagonism of the CCR9 receptor. Nature 450, 462–465 (2016).

56. K. Liu, L. Wu, S. Yuan, M. Wu, Y. Xu, Q. Sun, S. Li, S. Zhao, T. Hua, Z. Liu, Structural basis of CXC chemokine receptor 2 activation and signalling. Nature 585, 135–140 (2020).

57. J. A. Lees, J. M. Dias, F. Rajamohan, J. Fortin, R. O. Connor, J. X. Kong, E. A. G. Hughes, E. L. Fisher, J. B. Tuttle, G. Lovett, B. L. Kormos, R. J. Unwalla, L. Zhang, A. D. Schmitt, D. Zhou, M. Moran, K. A. Stevens, K. F. Fennell, A. E. Varghese, A. Maxwell, E. E. Cote, Y. Zhang, An inverse agonist of orphan receptor GPR61 acts by a G protein-competitive allosteric mechanism. Nat. Commun. 14 (2023).

58. X. Liu, S. Ahn, W. Alem, A. J. Venkatakrishnan, A. Masoudi, W. I. Weis, R. O. Dror, X. Chen, R. J. Lefkowitz, B. K. Kobilka, Mechanism of intracellular allosteric β2AR antagonist revealed by X-ray crystal structure. Nature 548, 480–484 (2017).

59. J. Duan, H. Liu, F. Zhao, Q. Yuan, Y. Ji, X. Cai, X. He, X. Li, J. Li, K. Wu, T. Gao, S. Zhu, S. Lin, M. W. Wang, X. Cheng, W. Yin, Y. Jiang, D. Yang, H. E. Xu, GPCR activation and GRK2 assembly by a biased intracellular agonist. Nature 620, 676–681 (2023).

60. K. Kobayashi, K. Kawakami, T. Kusakizako, A. Tomita, M. Nishimura, K. Sawada, H. H. Okamoto, S. Hiratsuka, G. Nakamura, R. Kuwabara, H. Noda, H. Muramatsu, M. Shimizu, T. Taguchi, A. Inoue, T. Murata, O. Nureki, Class B1 GPCR activation by an intracellular agonist. Nature 618, 1085–1093 (2023).

61. A. Jazayeri, R. M. Cooke, K. Hollenstein, J. Kean, A. Bortolato, R. K. Y. Cheng, A. S. Dore, M. Weir, F. H. Marshall, Structure of class B GPCR corticotropin-releasing factor receptor 1. Nature 199, 438–443 (2013).

62. Y. Q. Ping, C. Mao, P. Xiao, R. J. Zhao, Y. Jiang, Z. Yang, W. T. An, D. D. Shen, F. Yang, H. Zhang, C. Qu, Q. Shen, C. Tian, Z. jian Li, S. Li, G. Y. Wang, X. Tao, X. Wen, Y. N. Zhong, J. Yang, F. Yi, X. Yu, H. E. Xu, Y. Zhang, J. P. Sun, Structures of the Glucocorticoid-Bound Adhesion Receptor GPR97–Go Complex (Springer US, 2021; 10.1038/s41586-020-03083-w)vol. 589.

